# Blood flow synchronization in renal microcirculation - a high-resolution imaging study

**DOI:** 10.1101/2021.10.15.464509

**Authors:** Dmitry D. Postnov, Donald J. Marsh, Will A. Cupples, Niels-Henrik Holstein-Rathlou, Olga Sosnovtseva

## Abstract

**Aims:** Internephron signalling and interaction are fundamental for kidney function. Earlier studies have shown that nephrons signal to each other over short distances and adjust their activity accordingly. Micropuncture experiments revealed synchronous clusters of 2-3 nephrons formed from such interactions, while imaging and modelling results suggested the possibility of larger clusters. Such clusters are expected to play an important role in renal autoregulation, but their presence has not been confirmed and their size has not been estimated. In this study, we present methodology for high resolution renal blood flow imaging and apply it to estimate frequency and phase angle differences in kidney blood vessels under normal conditions and after administration of the vasoactive agents angiotensin II and acetylcholine.

**Methods and results:** To resolve signals from separate arterioles in a sufficiently large field of view, we developed a method for renal laser speckle contrast imaging. Our setup provides imaging of blood flow in the kidney cortex with a limit of image resolution at 0.8*μ*m per pixel and imaging frequency of 160Hz. We used the method to record from 1.5×1.5 mm^2^ sections of the renal surface in anaesthetised Sprague-Dawley rats in unstimulated conditions and during IV infusion of the vasoconstrictor angiotensin II or the vasodilator acetylcholine. In each section, we resolved and segmented 94.8±15.66 individual arterioles and venules, and analyzed blood flow using wavelet spectral analysis to identify clusters of synchronized blood vessels.

**Conclusions:** We observed spatial and temporal evolution of blood vessel clusters of various sizes, including the formation of large (>90 vessels) long-lived clusters (>10 periods) locked at the frequency of the tubular glomerular feedback (TGF) mechanism. The analysis showed that synchronization patterns and thus the co-operative dynamics of nephrons change significantly when either of the vasoactive agents is administered. On average, synchronization was stronger (larger clusters, longer duration) with angiotensin II administration than in the unstimulated state or with acetyl choline. While it weakens with distance, increased synchronization duration spanned the whole field of view, and likely, beyond it. Neighbouring vessels tend to demonstrate in-phase synchronization, especially in the vasoconstricted condition, which is expected to cause locally increased pressure variation. Our results confirm both the presence of the local synchronization in the renal microcirculatory blood flow and the fact that it changes depending on the condition of the vascular network and the blood pressure, which might have further implications for the role of such synchronization in pathologies development.

## 1 Introduction

The kidney represents a unique demand-driven, interconnected resource distribution network that is responsible for body homeostasis maintenance over a broad range of conditions, including variations in blood pressure and fluid intake and loss. With blood serving as the resource, single-nephron autoregulation mechanisms provide and regulate the demand. Based on measurements of tubule pressure responses to step changes in arterial pressure^1^, vascular transfer functions^2–7^, and renal blood flow response to arterial pressure forcing^8^, two critical mechanisms in renal pressure autoregulation, the myogenic mechanism and tubuloglomerular feedback (TGF), have emerged as the critical components. Their actions combine to regulate blood flow, serving to maintain the delivery of water and solutes to various regions of the nephron at levels appropriate to their dynamic ranges. The two mechanisms operate at different time scales, generating spontaneous blood flow and pressure oscillations at different frequencies: 5-10 seconds (0.1-0.2 Hz) for the myogenic response and 30-50 seconds (0.02-0.033 Hz) for the TGF^4, 9–12^.

Nephrons, however, do not operate as stand-alone units. Within a single kidney all nephrons are linked via the renal vascular tree, which provides connections of different proximity - from few hundred microns for neighbouring nephrons separated only by their respective afferent arterioles; to nephrons only connected at the level of the renal artery, which plays a role of a single supply source for all the nephrons. Nephrons nested in such a network are bound to communicate and affect each other to some degree. In addition to interaction through the blood flow and pressure, nephrons were found to communicate via electrical signalling. Such interactions are proven to play a critical role in kidney function and can lead to complex co-operative dynamics and synchronization between nephrons.

Micropuncture experiments^13–16^ showed that neighbouring nephrons (originating from a common artery) adjust their TGF-mediated tubular pressure oscillations to attain a synchronised regime. Although these experiments confirmed the existence of synchronisation, only pairs or triplets of nephrons could be sampled at any one time. Assessment of cooperative efforts of a larger number of nephrons required a different approach. To address this challenge, several groups adopted laser Speckle Contrast Imaging (LSCI)^17^. In LSCI, media with moving light scattering particles, e.g. red blood cells, are illuminated with a near-infrared laser. The backscattered light, recorded by a camera, forms an interference pattern, which appears more or less blurred depending on the speed of the particles. This pattern is then analysed to obtain qualitative maps of particle velocity and thus a blood flow estimate^18,19^. First applied by Holstein-Rathlou et al. to map TGF oscillations over a large field of view, this method was later used to explore periodic activity in the myogenic frequency band^20^, analyse spatial correlations in the renal blood flow^21^, and study intra-renal drug distribution^22,23^. Although these results encouraged the large-scale synchronisation hypothesis, the method lacked resolution, both spatial and temporal, as well as signal-to-noise ratio, to confirm it convincingly.

In this paper, we further advance renal blood flow imaging methodology and confirm the presence of synchronised clusters spanning multiple nephrons for the first time at the level of individual arterioles and venules. Our LSCI setup and data processing approach allow imaging renal microcirculation with at 0.8 *μ*m per pixel spatial resolution and imaging frequency to 160 Hz for 1024×1024 pixels. We apply it to study synchronous cluster formation.

## 2 Methods

### 2.1 Animal preparation

All experimental protocols were approved by the Danish National Animal Experiments Inspectorate and were conducted according to the American Physiological Society guidelines. Male Sprague Dawley rats (Taconic, Denmark) with average weight ≈ 290 g (n=5) were used. Before starting surgical procedures, animals were anaesthetized in a chamber with 8% sevoflurane. During the surgery, sevoflurane concentration was reduced to a final concentration of ≈ 2%. Two catheters were inserted in the right jugular vein to allow continuous systemic infusion of drugs and saline. Another catheter was inserted in the carotid artery to measure mean arterial pressure with a pressure transducer (Statham P23-dB, Gould, Oxnard, CA). Then tracheotomy was performed, after which the rat was placed on a servo-controlled heating table maintaining body temperature at 37 °C and connected to a mechanical animal ventilator (60 breaths/min; 8 ml/kg bodyweight). To avoid secondary heartbeat and breathing artefacts, Nimbex (muscle relaxant, Sigma) was administered in a concentration of 0.85 mg/ml, first as a bolus injection of 0.5 ml, followed by a continuous intravenous infusion at a rate of 20 *μ*l/min. The left kidney was then exposed, and the left ureter was catheterized to ensure free urine flow. To reduce motion artifacts and avoid drying the kidney surface during the experiment, we placed the kidney in a plastic fixation holder, covered it with warm agarose solution (1% Agarose, Sigma, 99% saline) and put a thin (0.1mm) cover glass on top of the kidney. Metal thread (40 micrometres in diameter) was bent in a “U” shape and positioned on top of the cover glass at the flattest location of the kidney surface, marking the region of interest for imaging procedures. Following the surgical procedures, the animal was left to stabilize for 20 minutes. Experiments were continued only if the mean arterial pressure remained within 100–120 mmHg during the control period. At the end of the experiment, animals were euthanized by overdose of sevoflurane, followed by cervical dislocation.

### 2.2 Laser Speckle Contrast Imaging data acquisition

To assess microcirculation in the kidney cortex, we built a high resolution laser speckle contrast imaging setup. A single-mode fibre-coupled laser diode (785nm, LP785-SF100, Thorlabs, USA) controlled with a laser driver, and temperature controller (CLD1011LP, Thorlabs, USA) was used to deliver coherent light onto kidney surface with a power density of approximately 10mW/cm^2^, providing an optimal signal to noise ratio^24^. Backscattered light was collected by a zoom imaging lens (VZM 1000i, Edmund Optics) at 5x magnification and recorded with a CMOS camera (Basler acA2000-165umNIR, 2048×1088 pixels, 5.5 *μ*m pixel size) at an exposure time of 5 ms. A subset of 1024×1024 pixels was used for recording, from a 1.5×1.5mm region of the renal surface. In addition, a linear polarizing filter was placed in front of the objective to reduce artefacts from reflected light. While the imaging set up in this configuration allows imaging resolution1 of ≈ 0.8*μ*m/px at the frame-rate of 160 frames per second (fps), in our experiments, we found it optimal, in terms of the field-of-view and data storage, to acquire images at 50 fps and resolution of ≈ 1.5*μ*m/px.

Imaging was performed in a non-stimulated state (control) and following administration of angiotensin II (AngII), and acetylcholine (ACh), respectively. After collecting 20 min (2400 frames) of the baseline data, we initiated continuous administration of AngII at a concentration of 4 ng/ml and an infusion rate of 20 *μ*l/min to cause systemic vasoconstriction. The infusion lasted for 30 min, out of which the first 10 min were allocated for blood flow to stabilize, and the following 20 minutes were recorded. Fifteen min after completion of the AngII infusion, we infused ACh at a concentration of 0.0375mg/ml and a rate of 20 *μ*l/min, first permitting 10 minutes for blood flow to stabilize, and then recorded subsequent 20 minutes. Across all experiments, average arterial pressure was 112±2, 127±6 and 100±4 during the control, vasoconstrited (AngII) and vasodilated (ACh) conditions respectively.

### 2.3 Data analysis

#### Image registration

To allow high-resolution laser speckle contrast imaging of the renal microcirculation, we needed to reduce motion artefacts, as any lateral motion larger than 5-10 micrometres will prevent accurate estimation of the blood flow and further segmentation of microcirculatory vessels. Unlike brain imaging, when working with the kidney, there is no bone tissue that can be fixed to reduce respiratory motion, and applying even slight pressure on the kidney might result in abnormal blood flow due to the occlusion of the small vessels on its surface. At the same time, raw laser speckle images, or contrast images without temporal averaging, are not suitable for automated registration due to the absence of clear intensity landmarks^25^. To resolve this issue, we placed a “U” shaped metal marker on the cover glass, which moves along with the kidney, as described above. As the first step of analyzing the data, the marker is segmented in all frames via thresholding and then used to estimate the translation type geometrical transformation required to register images. Estimated geometrical transformation is then applied to the raw laser speckle images prior to performing the contrast analysis.

#### Contrast analysis

Registered laser speckle images were processed to calculate temporal contrast 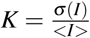, where *σ* (*I*) and < *I* > are standard deviation and mean of pixel intensity over 25 frames^25^. Contrast values were then converted to the blood flow index as *BFI* = 1*/K*^2^, which are then used in the ensuing analysis.

#### Vessels segmentation

To segment individual microcirculatory vessels, we calculated averaged in time BFI images and applied adaptive thresholding (MATLAB) to them. Automated segmentation was followed by manual clean-up, where we removed artefacts and occasional large surface vessels. We then calculated blood flow dynamics for each segmented individual microcirculatory vessel by averaging BFI values in pixels belonging to this vessel.

#### Synchronization analysis

To study synchronization patterns between microcirculatory vessels and, thus, obtain insight into inter-nephron communication, we apply continuous wavelet transform analysis (Morse wavelet, MATLAB) to segmented vessels’ BFI. We identified the frequency and phase of dominant periodic activity in the 0.015-0.05Hz frequency band associated with the TGF mechanism. Vessels with less than 10% prominence of the activity peak were discarded and not used for synchronization analysis. In this study, we consider blood flow in different segmented vessels to be synchronized whenever their dominant frequencies match. To quantify blood flow synchronization over the field of view, we analyzed phase differences between synchronized vessels, average synchronization duration and its dependency on the distance between vessels, and the probability of the vessel’s blood flow to be synchronized with N% of the vessels in the field of view. To provide a “single-value” characterization of the synchronization at a given moment of time, we also introduced the synchronization degree parameter *S*:

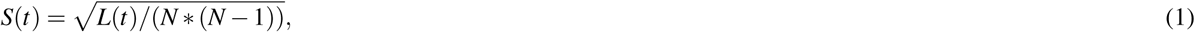

where *L* is a number of frequency matching pairs of vessels, *N* is a total number of observed segmented vessels, and *t* is the time. *S* represents the relation of the observed number of frequency matching pairs to their maximum possible number. Thus *S* = 1 corresponds to all segmented blood vessels having the same dominant frequency, while *S* = 0 to all segmented blood vessels having a different dominant frequencies. However, it is important to notice that *S* = 0 is impossible to reach due to the discrete nature of the measured data and the analysis. In our case, the 0.015 to 0.05 Hz range is split into 18 fixed values, so that if there were more than 18 vessels, it became unavoidable for some of them to have an identical dominant frequencies. In practice, for 100 vessels with randomly chosen dominant frequencies, the minimum observed *S* would be ≈ 0.25±0.06, which can be confirmed with a simple computational experiment.

### Statistical analysis

Paired t-test was applied to compare results between control, AngII infusion and ACh infusion. P-values greater than 0.05 are reported as not significant. Results were expressed as mean ± standard deviation (SD) unless indicated otherwise.

## 3 Results

### 3.1 Synchronization patterns

An example of a high-resolution blood flow map with segmented microcirculatory vessels is shown in Fig. 1,(c). Fig. 1,(d) shows RBF of four vessels outlined in (c) with corresponding colours (red, green, orange and black). From the corresponding dominant frequency and phase (Fig. 1,(e) and (f)), it can be seen that these vessels form two frequency-locked clusters (red-green and black-orange) that are synchronous most of the time. Moreover, there are long periods when these clusters synchronize, forming a larger cluster, interrupted with an asynchronous interval (from ≈ 850 to ≈ 1050 seconds).

**Figure 1.**
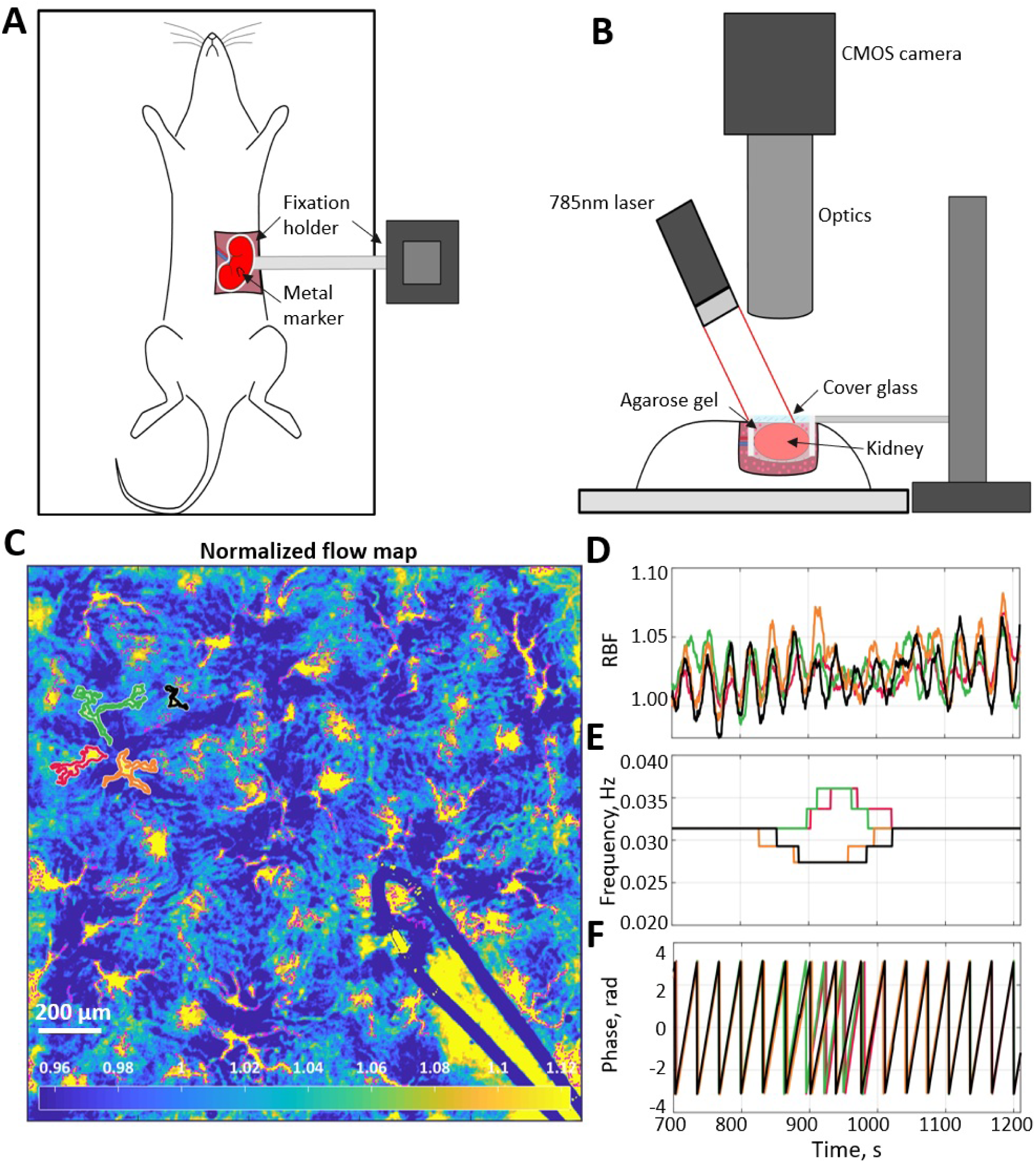
Schematics of the imaging setup and blood flow data example. (A),(B) Top and side view of the experimental setup and its key components. (C) Normalized flow map with microcirculatory vessels. Yellow and blue colours correspond to high and low blood flow, respectively. (D) Relative blood flow dynamics in four vessels, which are outlined in (C) with corresponding colours. (E) Dominant frequency in the TGF band of the blood flow oscillations is shown in (D). (F) Phase value at the dominant frequency. One can see that there are two vessels pairs in which blood flow is synchronized for more than 90% of time. Synchronization between the pairs is also well observed but breaks down when TGF activity change its frequency at 850–1050 s). These exemplary graphs reflect the dadata

We generalized this approach to the full-field blood flow imaging of the kidney surface. Figure 2 shows examples of high (A), moderate (B), and low (C) instantaneous synchronization degree, with *S* = 0.97, 0.65, and 0.23 respectively. In the first case, flow in almost all of the identified vessels oscillates at the same TGF frequency, forming a cluster of >100 vessels within the field of view. This cluster is likely to be even larger since it can spread outside the view field and in-depth in the kidney cortex. In the case of moderate synchronization degree - several frequency clusters with ≈ 10 *−* 60 vessels can be identified, while for the low *S* there are multiple groups of 2-3 vessels displaying the same frequency but no distinct pattern over the field of view. All three regimes were observed in the same animal during control, vasoconstricted (AngII infusion) and vasodilated (ACh infusion) conditions respectively. Averaged over the whole observation period and all animals (N=5, 20 min per condition) synchronization degree has moderate to high 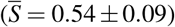 values during AngII infusion, low to moderate 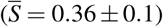 during ACh, and low to high in control 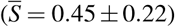.

**Figure 2.**
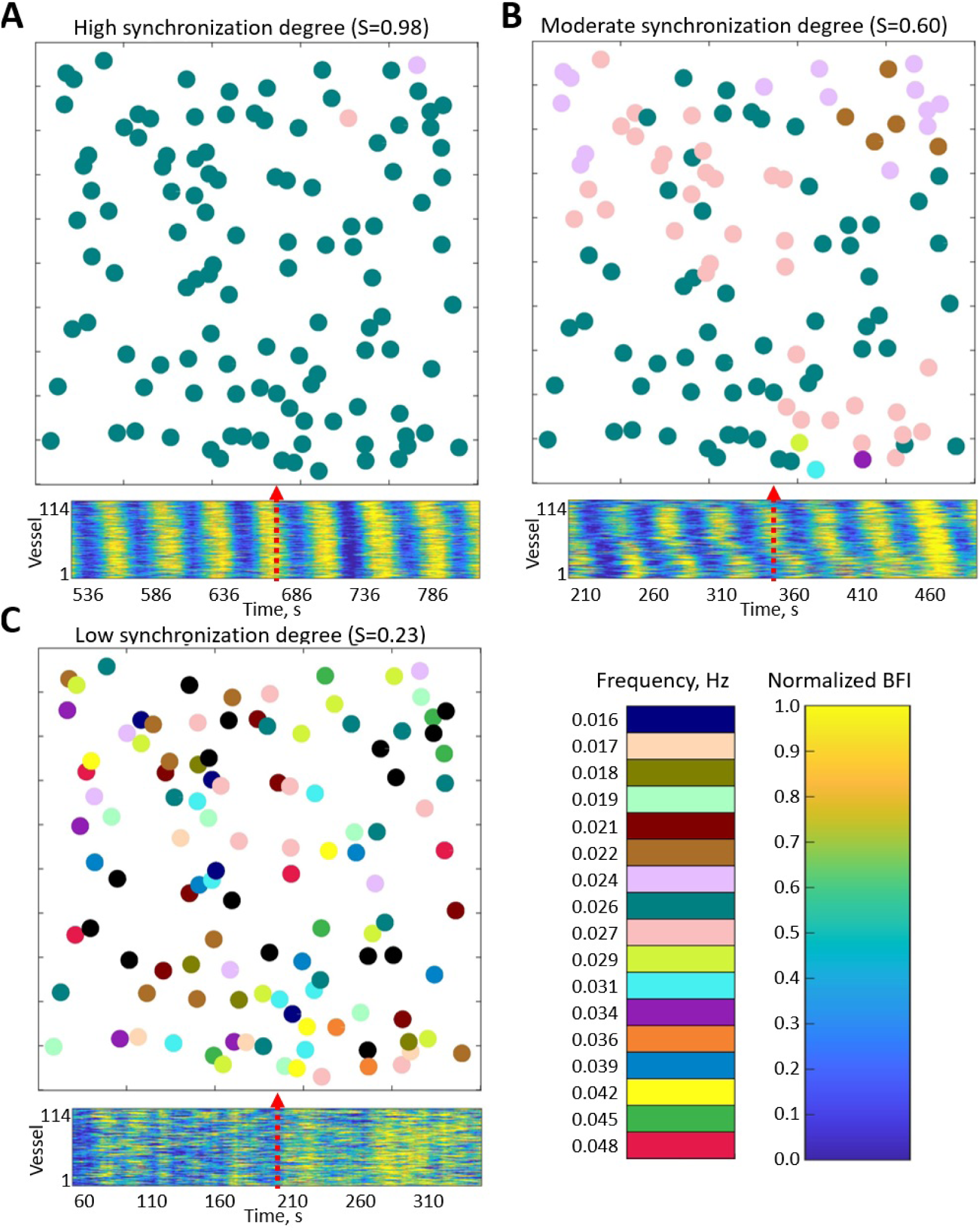
Examples of instantaneous frequency patterns. Each circle location corresponds to a segmented vessel with color-coded frequency. Bottom panels show normalized blood flow in each segmented vessel over 5 minutes centered around the time corresponding to the frequency pattern snapshot. (A) High synchronization degree *S* = 0.97 - large cluster covers the whole field of view. (B) Moderate synchronization degree *S* = 0.65 - most of the vessels are split between two clusters. (C) Low synchronization degree *S* = 0.23 - no clear clustering pattern, some vessels do not have pronounced TGF activity (frequency colour-coded as black). All data were captured in the same animal but under different conditions: control (A), AngII infusion (B), ACh infusion (C). Black-colored circles represent vessels where TGF activity was considered too weak (less than 10% prominence of the activity peak).

To visualize clustering tendencies, we calculate the probability of a randomly chosen vessel at any given moment of time to have the same dominant frequency as *X* % of the vessels in the field of view. From Fig. 3 it can be seen that during AngII infusion, vessels are significantly more likely to be synchronized with 20 − 50% of the field of view than in control or during ACh infusion. In the latter condition probability of a vessel being synchronized with less than 15% are significantly higher than during AngII infusion (p<0.05). Such behaviour is observed both with no restriction on minimum synchronization duration (A) and when only frequency-locking for 3 TGF periods and longer is taken into account.

**Figure 3.**
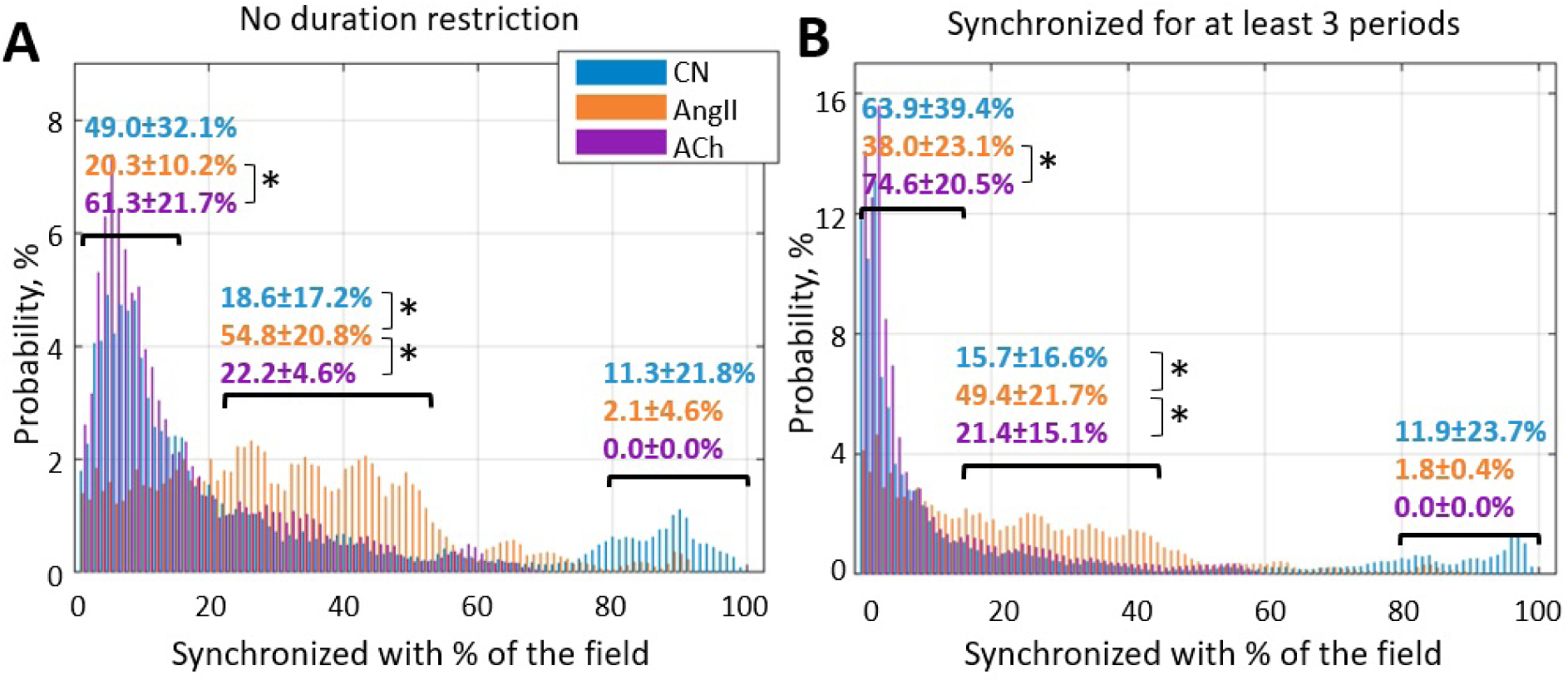
Probability of a randomly chosen vessel displaying the same dominant TGF frequency as *X* % of the vessels in the field of view. (A) Any duration of frequency matching is considered, and (B) dominant frequency should match for at least 3 TGF periods. It can be seen that infusion of AngII significantly (p<0.05) increases the prevalence of clusters covering 20-50% of the vessels - chances that a randomly chosen vessel belongs to such a cluster at any given moment are ≈ 50%, while for control and ACh infusion they are ≈ 20%.N=5 animals were used to create these graphs. Paired t-test was used to produce P-values. P-values smaller than 0.05 are considered to be significant and marked with “*”.

### 3.2 Phase waves and spatial localization

Another distinct feature that we have observed is that the phase of oscillations within a cluster is space-dependent - it is mostly the same for the closely positioned vessels and gradually changes with distance. Figure 4 illustrates how phase within clusters evolves in space and time, forming the phase waves. Note that both (A) and (B) panels are from the same animal as was shown in Fig. 3 in control and vasoconstricted conditions, respectively, and that only vessels belonging to the largest cluster are shown. Change of the wave direction by ≈ 90° in the same animal with constant vascular structure suggests different synchronization centres where phase waves originate. Propagation speed is ≈ 0.37 and 0.30 mm/s for (A) and (B) respectively and spatial period of the phase waves is ≈ 8.85 and 3.95 mm (respective phase difference |*δ*| ≈ 0.71 and 1.59 rad was observed over 1 mm distance). The possibility of such phase-waves was previously hinted at in our earlier laser speckle contrast imaging study, where spatial correlations in renal blood flow were analysed^21^. See supplementary videos for more examples of phase waves dynamics.

**Figure 4.**
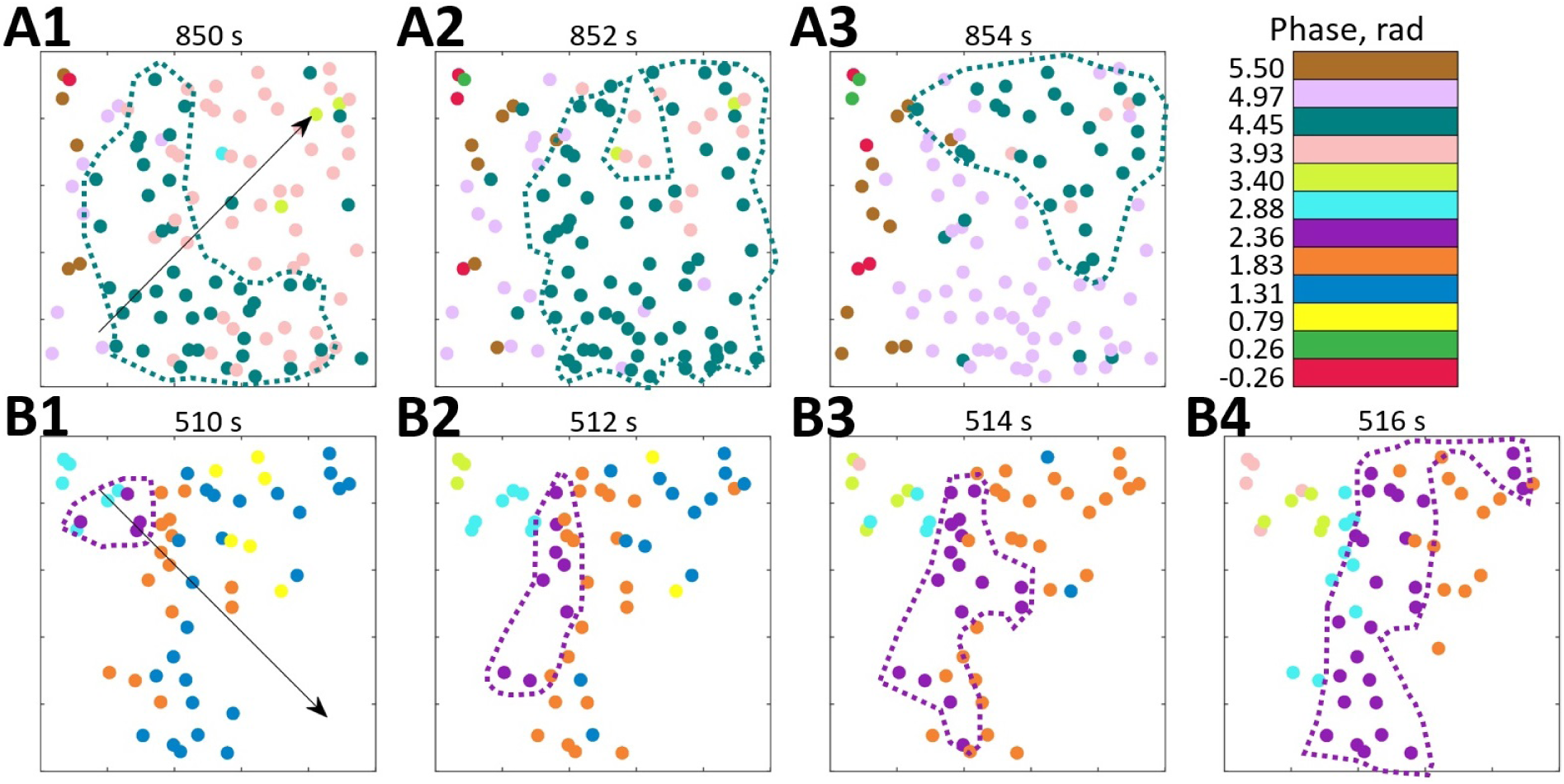
Phase waves. (A) and (B) show the spatio-temporal evolution of TGF activity phase in vessels that belong to the largest cluster observed during control and AngII infusion. Both observations are from the same animal as shown in Fig. 2(A) and (B). Arrows indicate the wave direction.

Phase difference distribution (Fig. 5, A), calculated over all of the frequency matching vessels in all animals, shows the prevalence of in-phase synchronization 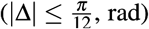. This result is in good agreement with experimental micropuncture observations, where it is explained by the presence of fast electrical coupling acting over short distances. In all conditions, the larger phase differences are less prevalent, with the anti-phase synchronization 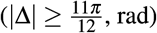 observed over just 1% of time in control and during ACh infusion. However, AngII infusion increases this number to 5%, showing a statistically significant difference with other conditions. Phase difference grows with distance, as can be seen from Fig. 5 (B), reflecting the presence of phase waves. While change is relatively small in the control and vasodilated conditions (≈ 0.3 and 0.25 rad/mm), it is strongly enhanced in the vasoconstricted condition, reaching, on average, ≈ 1 rad over 1 mm distance. As we showed with mathematical modelling^26^, such difference can be explained by strengthened hemodynamic coupling, which the increased vascular tone should cause.

**Figure 5.**
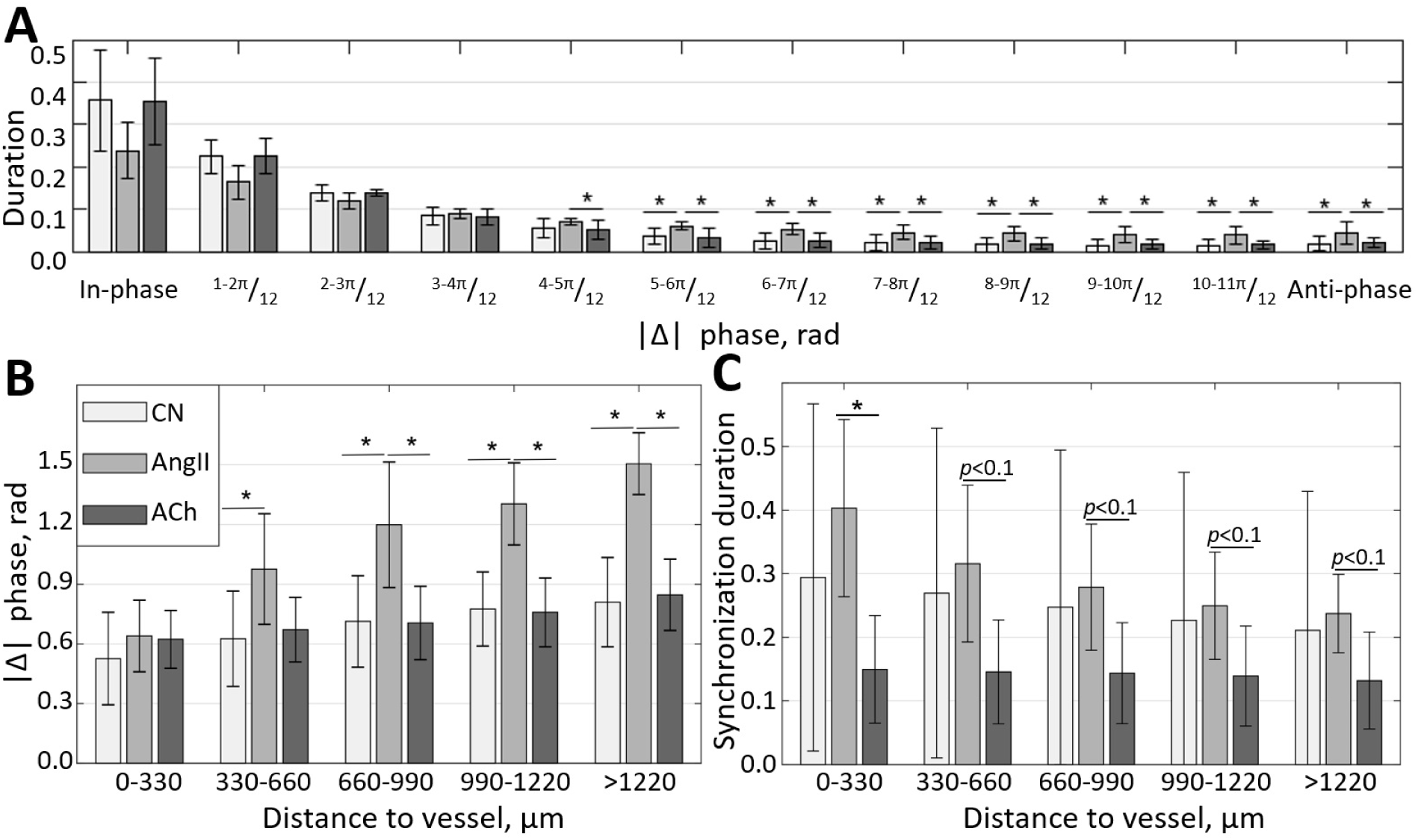
Localization of phase and synchronization in space. (A) Phase differences prevalence for synchronized vessels. (B) Phase differences distribution over the distance between vessels. (C) Synchronization duration normalized by the total observation time for different distances between vessels.N=5 animals were used to create these graphs. Paired t-test was used to produce P-values. P-values smaller than 0.05 are considered to be significant and marked with “*”. Values between 0.05 and 0.1 are shown as *p* < 0.1, where relevant, to highlight a trend in the data.

Since synchronization in renal blood flow is extended far beyond a pair of nephrons and is unlike synchronization in relatively homogeneous media, one would expect some space localization due to the topological features of the vascular network. Figure 5(C) illustrates how average synchronization duration changes with distance. It is clear that synchronization is stronger in the vasoconstricted condition, with average duration reaching ≈ 40±13% of the observation time for neighbouring vessels and gradually reducing to 24±6% for vessels located at more than 1 mm distance. In control, synchronization duration varies greatly, but on average, it also reduces with distance, although at a slower rate than during AngII infusion - from ≈ 30% to ≈ 21%. During the ACh infusion the synchronization duration is ≈ 15 *−* 13%, with only 2% reduction over 1 mm of distance. Higher synchronization duration in control and during AngII infusion compared to ACh infusion suggests a long-distance nephron-to-nephron communication or a common driving force.

## 4 Discussion

In this study, we have designed a methodology for high-resolution blood flow imaging in renal microcirculation and applied it to study the synchronization of TGF oscillations in control, vasoconstricted (AngII infusion) and vasodilated (ACh infusion) conditions. Our data confirm that blood flow in renal microcirculation tends to demonstrate clustered, frequency-locked activity, with the clustering size and tendencies changing depending on the animal condition. Synchronization tends to be stronger, and cluster size is larger during AngII infusion, with synchronization degree 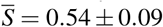 and average synchronization duration ranging from ≈ 40±13% to 24±6% of the observed time depending on the distance between vessels. During the ACh infusion, on the contrary, synchronization seems to be disrupted, with 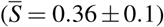 and average synchronization duration ≈ 15±7*−*13±7% of the observed time. Finally, in the normotensive condition, we observed mixed behaviour with highly variable synchronization degree 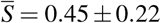 and duration ranging from ≈ 30±27% to 21±21%. A possible explanation for the stronger synchronization during the AngII infusion will be a stronger hemodynamic coupling due to increased vascular resistance. It is also supported by the increased number of vessels synchronized in anti-phase during the infusion, which, as we predicted using mathematical modelling^26^, is a natural consequence of stronger hemodynamic coupling.

We also observed phase waves that travel over the TGF-frequency clustered vessels (Figs 4, and 5(B)). Interestingly, the direction of the phase waves is not predetermined by the vascular structure - depending on the flow dynamics and other, yet unknown factors, it can change even within the same animal (see Fig. 4). Such behaviour might be related to diffusive interaction between nephrons or the formation of synchronization centres (pacemakers) at different locations. The presence of phase waves supports the deterministic nature of synchronization rather than random entrainment at the same dominant TGF frequency. Similar effects are well studied in excitable media with diffusion interaction mechanisms such as brain^27,28^ and heart^29^ tissues. Such similarity might suggest a presence of a diffusive mechanism in the inter-nephron interaction or a long-distance fast electrical signalling, e.g. via conducted vasoreactivity.

In our experiments, we imaged 1.5×1.5 *mm*^2^ field of view in 5 animals with 94.8±15.66 individual segmented vessels on average, each of which is 10 *−* 30*μ*m in diameter. While it provides an estimate of synchronization in the nephrons activity, direct translation from individual segmented vessels to nephrons is challenging and requires further exploration. Factors that are critical to consider are (i) penetration depth of LSCI when applied to renal imaging, (ii) type of the vessels in the field of view, and (iii) topology of the nephro-vascular network. While in theory, LSCI can collect the blood flow signal from as deep as 300-400 *μ*m, in practice, visually resolvable signal typically comes from top 50-150 *μ*m of the vascular structure^30,31^. Considering high vascular density close to the renal surface, it would mean that LSCI is likely limited to imaging vessels originating from ≈ 10000 nephrons in outer 30% of rat renal cortex^32^, which would result in ≈ 40 nephrons in the 1.5*x*1.5*μ*m field of view. The larger number of segmented vessels can be explained by their mixed type - afferent and efferent arterioles as well as venules are likely to be segmented. Distinguishing vessels types in LSCI images will require further exploration and registration with high-resolution structural imaging. It, however, does not mean that the observed clusters were limited to, at the most, 40 nephrons. When considering renal vascular topology, it is to be expected that within the 1.5*x*1.5*μ*m field of view, we observe arterioles that arise from different non-terminal arteries^33^. Depending on the branching order, each of such arteries can branch into ten-several hundreds of nephrons, but only a small number of these nephrons will have arterioles reaching close enough to the surface to be segmented from LSCI images. Thus, when a synchronous cluster is observed with LSCI, it is likely to extend several branching orders in depth and reach the size of hundreds and even thousands of nephrons.

While the exact role of inter-nephron communication, co-operative dynamics and synchronization in kidney-related pathology development is still unclear and requires further exploration, it is evidently altered by the blood pressure and vascular tone. Strong local coupling and in-phase synchronization, while being not evident at the renal artery level^22^, are likely to increase pressure variation at the level of afferent arterioles^26,34^, thus increasing chances of local damage and aggravating pathological condition.

## Data Availability Statement

The data underlying this article will be shared on reasonable request to the corresponding author.

